# Comparable human spatial memory distortions in physical, desktop virtual and immersive virtual environments

**DOI:** 10.1101/2022.01.11.475791

**Authors:** Fiona E. Zisch, Coco Newton, Antoine Coutrot, Maria Murcia, Anisa Motala, Jacob Greaves, William de Cothi, Anthony Steed, Nick Tyler, Stephen A. Gage, Hugo J. Spiers

**Author notes:** These authors contributed equally to this work (Fiona E. Zisch, Coco Newton, Antoine Coutrot).

## Abstract

Boundaries define regions of space and are integral to episodic memories. The impact of boundaries on spatial memory and neural representations of space has been extensively studied in freely-moving rodents. But less is known in humans and many prior studies have employed desktop virtual reality (VR) which lacks the body-based self-motion cues of the physical world, diminishing the potentially strong input from path integration to spatial memory. We replicated a desktop-VR study testing the impact of boundaries on spatial memory (Hartley et al., 2004) in a physical room (2.4m x 2.4m, 2m tall) by having participants (N = 27) learn the location of a circular stool and then after a short delay replace it where they thought they had found it. During the delay, the wall boundaries were either expanded or contracted. We compared performance to groups of participants undergoing the same procedure in a laser-scanned replica in both desktop VR (N = 44) and freely-walking head mounted display (HMD) VR (N = 39) environments. Performance was measured as goodness of fit between the spatial distributions of group responses and seven modelled distributions that prioritised different metrics based on boundary geometry or walking paths to estimate the stool location. The best fitting model was a weighted linear combination of all the geometric spatial models, but an individual model derived from place cell firing in Hartley et al. 2004 also fit well. High levels of disorientation in all three environments prevented detailed analysis on the contribution of path integration. We found identical model fits across the three environments, though desktop VR and HMD-VR appeared more consistent in spatial distributions of group responses than the physical environment and displayed known variations in virtual depth perception. Thus, while human spatial representation appears differentially influenced by environmental boundaries, the influence is similar across virtual and physical environments. Despite differences in body-based cue availability, desktop and HMD-VR allow a good and interchangeable approximation for examining human spatial memory in small-scale physical environments.

## (1) Introduction

Returning to a room to replace a recently collected object one might plausibly retrace one’s steps, using knowledge of the self-motion on the path to the place where the object should be replaced. Alternatively, one could walk to the location in the room that matches the visual memory from the wall boundaries or a specific feature in the room to locate where the object should be replaced. It is also plausible that all these strategies could be combined. Numerous studies have indicated that humans and animals use such strategies to recall spatial locations and navigate (Ekstrom, Spiers, Bohbot, & Rosenbaum, 2018; Gallistel, 1990; O’Keefe & Nadel, 1978; Spiers & Barry, 2015). Returning to a location using processing of self-motion is referred to as path integration, whereas using visual information about the distance between walls or objects is referred to as a geometric strategy (Gallistel, 1990).

Much of our understanding of how geometry and path integration are used in spatial memory has come from studying ‘place cells’ in the rodent hippocampus, which express localised activity in the environment, providing a neural foundation for a cognitive map (O’Keefe & Nadel, 1978). Seminal studies in the 1990s revealed evidence for both path integration and visual geometry driving place cell firing (Gothard, Skaggs, & Mcnaughton, 1996; O’Keefe & Burgess, 1996). O’Keefe and Burgess (1996) found that expansion and contraction of wall boundaries of an environment rats were foraging in had predictable stretching and compression of the location-specific activity of place cells, indicating that they were driven by the distance and allocentric direction to local wall boundaries. This led to the development of models seeking to explain the activity patterns as a set of boundary vector inputs (Burgess & O’Keefe, 1996; Grieves, Duvelle, & Dudchenko, 2018; Hartley, Burgess, Lever, Cacucci, & O’Keefe, 2000). Such boundary vector codes have since been observed in the activity of the subiculum in rodents (Lever, Burton, Jeewajee, O’Keefe, & Burgess, 2009) and humans (Lee et al., 2018; Stangl et al., 2021). These are functionally relevant – rats with a lesion to hippocampus were impaired in finding goals based on environment shape cues but not visual features (McGregor, Hayward, Pearce, & Good, 2004). Further evidence for how boundaries impact on the neural representations of space has come from the study of ‘grid cells’ in the medial entorhinal cortex, which show periodic repeating patterns of activity forming a tessellating array across the space of the environment. Compression and expansion of wall boundaries has been found to cause grid cell patterns to compress or expand in relation to the amount of change (Barry, Hayman, Burgess, & Jeffery, 2007) and distort in a predictable manner when environments are transformed from rectangular to trapezoidal shapes (Krupic, Bauza, Burton, Barry, & O’Keefe, 2015). Grid field locations and orientation are also highly dependent on environment symmetry (Stensola, Stensola, Moser, & Moser, 2015). Thus, rodent experiments manipulating the size and shape of environments has provided useful insights into how the brain represents space.

Given that place and grid cells are thought to underlie our long-term spatial memory and their activity patterns change in a predictable manner following expansions and contractions of the environments, a logical corollary is that human spatial memory should follow similar patterns of updating after an expansion or contraction of that environment. This assumption was tested by Hartley et al. who had humans learn the location of an object and subsequently replace it in varying sized rectangular or square environments rendered in a desktop virtual reality (VR) environment (Hartley, Trinkler, & Burgess, 2004). The resulting distributions of the replaced object mimicked the characteristic stretched and compressed pattern of place cell firing and were best predicted by a model based on place cell activity relating the change in the distance to local walls in the environments. This ‘boundary proximity’ model predicted object replacements to maintain fixed distances to nearby walls for objects found near boundaries or during room expansions, and fixed ratios of distances between opposing walls for more central objects or during room contractions. This was true even for disorientated responses, when participants appeared to apply geometric spatial metrics to the wrong axes of the environment, suggesting an independence of location and orientation estimation processes (Hartley et al., 2004). Indeed, other studies using immersive head-mounted-display (HMD) VR have also highlighted the influence of boundary geometry for spatial orientation (Kelly, Mcnamara, Bodenheimer, Carr, & Rieser, 2009). Spatial memory tasks in HMD-VR enable use of body-based self-motion cues. Participants either walk across the room or can walk on a static foot-tracking frame that allows walking patterns to simulate movement (Diersch & Wolbers, 2019). These experiments have also provided evidence that changes in spatial memory after non-orthogonal (e.g. trapezoidal) changes to the wall boundary layouts can additionally be explained by models using the expansion and contraction observed in grid cell activity (Bellmund et al., 2020)

While VR has significantly advanced our understanding of human spatial memory, it remains unclear how the influence of boundaries replicates across virtual and physical environments (Montello, Hegarty, Richardson, & Waller, 2004). Manipulating boundaries in physical spaces presents a host of practical problems that are solved by VR, such as instantaneous environmental manipulation without wear and tear of materials that can introduce additional spatial cues that are hard to control for. Desktop-VR specifically is compatible with functional magnetic resonance imaging (fMRI), within which virtual movement is controlled by a joystick or arrow keys and self-motion information is derived from optic flow only (Ekstrom et al., 2018; Epstein, Patai, Julian, & Spiers, 2017; Spiers & Barry, 2015). Conversely, HMD-VR purportedly offers a higher degree of environmental presence and allows for body-based self-motion cues such as vestibular and proprioception feedback (Klatzky, Loomis, Beall, Chance, & Golledge, 1998; Sousa Santos et al., 2009), but see (Feng, Duives, & Hoogendoorn, 2022). Both have been critical in elucidating the brain-behaviour relationships of navigation (Keinath, Rechnitz, Balasubramanian, & Epstein, 2021; Shine, Valdés-Herrera, Hegarty, & Wolbers, 2016). However, the specific advantages of different VR environments to explore boundary manipulation are only relevant if they faithfully recapture the spatial memory processes used by humans in physical environments. A large body of work has already addressed this knowledge gap, comparing different aspects of spatial memory such as the influence of environmental deformations, visual cue conflict and wayfinding strategy across either one VR type and physical environments, or between desktop and immersive-VR environments (Feng et al., 2022; Keinath et al., 2021; Kimura et al., 2017). However, few studies have compared the impact of boundaries on human spatial memory across all three.

Given the dependence of neural firing and spatial behaviour on boundaries, we sought here to compare how expansion and compression of square and rectangular wall boundaries might impact human spatial memory of an object’s location in physical space, desktop-VR and HMD-VR. We were able to leverage the use of the engineering facility PAMELA (Pedestrian Accessibility Movement Environment Laboratory) to build a uniform, four-walled room with manually moveable walls, which we could then replicate virtually using high-precision laser scanning. Following the original Hartley et al (2004) experimental paradigm, we considered a range of geometric models that might predict human memory for the object’s location, using either simple distance and angular spatial metrics or place cell firing activity. Based on evidence from Hartley et al. (2004) we predicted response distributions would be best accounted for by models using a fixed distance or fixed ratio to the walls depending on the object location and if the room was expanded or contracted (i.e. the boundary proximity model formulated in Hartley et al, 2004). We then aimed to explore the contribution of body-based self-motion cues by examining whether HMD-VR response distributions and predictive model fits would be more similar to the physical space or desktop-VR, and whether additional path-integration based models could effectively capture response distributions across the environments.

## (2) Methods

To test how different spatial distortions affect human memory and how this might differ between a physical and a virtual environment, we ran three quasi-identical experiments investigating memory of object location and the impact of spatial geometric distortion thereupon in a) a physical ‘real-world’ environment (Physical), b) an immersive virtual reality environment (HMD-VR), and c) a non-immersive virtual reality environment (Desktop-VR). Each of these saw us change the environmental boundaries twice; first, the learning environment was changed from an initial “small” setup (2.4m x 2.4m) to the testing environment’s “large” setup (4.8m x 4.8m). The learning stage object was placed at 1.95m x 0.3m relative to the 0,0 position of the Southwest corner. Second, the learning environment was changed from a “large” setup (4.8m x 4.8m) to a “tall” testing environment (2.4m x 4.8m) and the learning object placed at 2.9m x 2.9m relative to the 0,0 position.

In all environments and experimental phases, we used object position tracking before and after spatial changes had been made to calculate which models were used and then compared these both across participants and across trials. All participants gave informed, signed consent to participate and were compensated £10 for the experiment. Ethics approval was given by the UCL ethics committee (Project Number: CPB/2013/015).

### (2.1) Experiment 1 - Physical environment

#### (2.1.1) Methods

##### (2.1.1.1) Materials

A four-walled testing environment was erected on a hydraulic platform at the UCL PAMELA research facility. The walls of the enclosure were made of four plywood panels, each wall 4.8m wide x 2.4m tall (See Figure and Appendix). The panels were painted white and had furniture gliders attached to their base in order to be able to move walls easily on an anthracite carpet we installed on the platform. Given the difficulty of manually moving the four walls in the physical PAMELA environment, only two transformation types could be implemented. Furthermore, given the time required to physically move the walls and group testing of participants, trials were always completed by participants in the same order and were not randomised. This differs from Hartley et al 2004, where 16 transformations were used in different orders. However, from our pilot experiment, we found two trial types were sufficient to dissociate model fits from predictions of the different models. The three configurations used here were selected from the configurations tested in the pilot study and the transformation types designed to optimally test predictions from the Boundary Proximity Model. The ‘large’ room configuration was 4.8m x 4.8m; the ‘tall’ room was 2.4m (wide) x 4.8m (tall); and the ‘small’ room was 2.4m x 2.4m.

It was necessary to provide a way for participants to orient themselves globally, as they entered the environment from a different corner in each phase while blindfolded (comparable to the procedure of teleporting in desktop VR followed in Hartley et al, 2004). Hartley et al.’s experimental design used a mountain range projected at infinity as a directional cue. In a physical setup this mode of operation is not feasible. Projecting a panorama onto the wall would provide a landmark rather than distal cue, which would result in an environmental feature rather than a constant, allowing participants to orientate within the environment. We used PAMELA's programmable LED lighting to light one side of the room in amber as an orientation cue. The row of amber lights remained constant throughout the experiment and provides a non-geometric and non-landmark cue. A blackout curtain was suspended from above, obstructing views of the surrounding cues within the research facility.

The object used for both learning and recall was a stool (white Tam Tam plastic stools from Habitat). The stool is easy to hold even when blindfolded and provides no misleading (or otherwise) orientation cues to the participant. It was chosen due to its specific rotational symmetry along the vertical axis, eliminating the question of object orientation. The stool has a diameter of 31cm at the widest section and a height of 45cm.

Twelve ‘Optitrack’ cameras were used to track the final location of the stool. To enable this to work properly, there were three retro-reflective markers on the top of the object. The cameras tracked the location of both the object and the participant (wearing a hat with markers), both reflecting infrared light. The data was recorded using the software ‘Motive’. As a backup to the tracking system failing - as evidence in the pilot study - a video camera was placed above the centre of the room so that the final object location could be estimated from video footage.

##### (2.1.1.2) Participants

29 participants (18 female, 11 male; average age 40.03/44.8 years, SD = 15.47/15.7) were recruited from the student and staff population at University College London. One participant was older than the required age range (18-65) and was excluded from the results. Another participant was partially sighted and therefore was also excluded.

##### (2.1.1.3) Procedure

In each session of the study, groups of participants (between 6 and 9) gathered in the waiting room for an induction. An experimenter explained the task and answered questions. Instructions were to memorise the position of the stool, which they would then have to replace at a later stage. Participants were encouraged to perform as quickly and accurately as possible and were not informed about potential manipulations to the room geometry or different starting locations per trial at any point.

To replicate the teleporting used by Hartley et al (2004) in the physical environment and to ensure that participants could not use spatial information from outside the testing enclosure to situate and orient within an environmental reference frame, participants were blindfolded before leaving the waiting area and led into and out of the testing area by an experimenter. They were also disoriented before and after being in the testing room to ensure that they could not use self-motion cues to deduce reference information from outside the room.

Two transformations were used: small room configuration to large configuration (Trial 1) and large configuration to tall configuration (Trial 2), both involving a learning and placement stage (Table 1). In the learning stage, participants were led to the starting position in a corner of the room facing the centre. They removed their blindfold when a beep noise sounded. They retrieved the object, put the blindfold back on and notified the experimenter who then led them out of the room. In the waiting period, the testing room’s size and shape were changed; participants were prevented from interacting with each other to maintain independency of results. After a short pause in a waiting area, participants re-entered the room similarly blindfolded but at a different corner starting position and facing the walls while holding the object. They replaced the object where they remembered it to have been before, and again left the room guided by an experimenter. Starting positions were the same across all participants, unlike the main experiment of Hartley et al.

**Table 1.** Stages for experimental trials 1 and 2 in pilot study and experiments 1, 2, 3.

| **Trial** | **Experimental stage** | **Task** |
| --- | --- | --- |
| Trial 1 | Learn 1 | Retrieve the object and memorise location 1 |
|  | Place 1 | Place the object in the location memorised in learning stage 1 |
| Trial 2 | Learn 2 | Retrieve the object and memorise location 2 |
|  | Place 2 | Place the object in the location memorised in learning stage 2 |

Participants completed three experimental trials: a practice trial where the room was not manipulated between learn/place stages, followed by two testing trials with room manipulations. The practice phase was conducted in the large 4.8m x 4.8m room configuration. After all trials were completed, experimenters debriefed participants about their experience and strategy of remembering the stool location. The experimental task lasted approximately 1.5 hours.

### (2.2) Experiment 2 - Head-mounted display immersive virtual environment

#### (2.2.1) Methods

##### (2.2.1.1) Materials

The virtual environment consisted of high-fidelity point clouds obtained from 3D laser scans of the physical environments, rendered with a bespoke GPU-based point cloud renderer. 3D scanning was performed with a Faro Focus X330 laser scanner. The scanned floor was substituted by a texturised plane to circumvent the otherwise missing points resulting during scanning from the scanner's blind spot beneath it. The VEs were rendered in a HTC Vive Developer Edition at 1:1 scale in Unity at 90FPS with a vertical FOV of 60 degrees. The HTC Vive Developer Edition base stations were placed in opposite corners of the 4.8m x 4.8m surface area (corresponding to the largest room configuration).

Participants used one HTC Vive Developer Edition wireless controller throughout the study. In each trial, the application would always begin with an empty virtual space where only the virtual controller was visible. Participants could then begin the corresponding stage of the experiment by pressing the front red system button. During each stage, participants could grab the object by pressing the rear trigger whenever the controller was within a 5cm range of the object. By releasing the trigger button, the object would then fall to the floor level and rotate to a vertical position. The object used in the study was a virtual replica of the white Tam Tam plastic stool.

##### (2.2.1.2) Participants

39 participants (14 female, 25 male; average age 30.8 years, SD = 10.9) were recruited from the student and staff population at UCL.

##### (2.2.1.3) Procedure

The virtual room, trial configurations and instructions were again identical to the physical setup. The experimental protocol was adapted for the VR trials to match the physical trials in experiment 1 as closely as possible. The study was conducted in two different rooms at University College London: a waiting room and a VR testing room.

Participants were informed that they would be able to see the SteamVR chaperone grid when using the VR system. They were told that this meant they were approximately 0.4m away from the physical limits of the room and asked to ignore it when completing the experimental task. They were also shown a physical version of the Tam Tam white stool from Habitat. Prior to the two experimental trials, participants completed a practice trial. This was to ensure that they had a chance to familiarise themselves with the HTC Vive controller, as well as grabbing and placing the virtual object. The practice VE consisted of an empty space with a floor and a sphere (radius=0.25m) to practice interacting with the controller. Participants were asked to navigate to the sphere. They were then asked to grab and release it as many times as needed until they felt comfortable with the interaction.

As in experiment 1, the experiment consisted of two trials and each trial involved a learning stage and a placing stage.

The experimenter disoriented and guided the participant to the starting location for the corresponding stage (using the HMD as a blindfold). Participants were disoriented to stop them from finding correspondences between the physical testing room and the VE. Starting locations were marked with tape in the testing room to help the experimenters place participants at the correct starting location and facing direction for each stage.

### (2.3) Experiment 3 - Desktop non-immersive virtual environment

#### (2.3.1) Methods

##### (2.3.1.1) Materials

The immersive VR environment used in experiment 2 was reprogrammed into a desktop navigable environment for experiment 3. Participants used a mouse and keyboard to navigate the desktop environment; keyboard arrow keys allowed participants to move forward and backward, left and right, movement of the mouse provided means to alter their viewpoint on a 360° axis around themselves, so that movement and gazing in all directions was possible. Left clicking the mouse would pick up the stool when standing within 5cm of it, and right clicking dropped it in front of them. The computer for the task was situated on a desk within a small testing room (2m x 4m x 3m). In order to replicate a feeling of isolation within the four-walled room, the room was darkened completely apart from the monitor screen. As the experiment was run in the evening, no light came through the windows and the blinds were drawn to remove external distractors.

##### (2.3.1.2) Participants

A total of 44 participants (24 female, 20 male; average age 27 years, SD = 9.0) were recruited from the UCL online psychology participant pool.

##### (2.3.1.3) Procedure

The virtual room, trial configurations and instructions were again identical to the physical setup, as well as the immersive VR setup. The experimental protocol was adapted for the desktop trials to match the physical and immersive VR trials in experiments 1 and 2 as closely as possible.

Participants had an opportunity to practice using the controls on a laptop in the waiting room. Participants had to pick up the stool and repeatedly move it between two black circles in the virtual room until comfortable with the controls.

Experiment 3 was conducted in two rooms at University College London, a waiting room and a testing room. Participants were led from the waiting room to the testing room by an experimenter where they first conducted trial one, followed by a break in the waiting room and then completed trial two.

### (2.4) Models

The geometric models considered are Absolute Distance (AbsDist), Corner Angle (CorAng), Fixed Ratio (FixRat), Fixed Distance (FixDist), and Boundary Proximity (BoProx) as a weighted combination of the previous two from Hartley et al. 2004. In addition, we considered Path Integration Fixed (PathIntFix) and Path Integration Ratio (PathIntRat), see Table 2 and Figure 1. Despite the original Hartley et al. 2004 task design aim to minimise contributions of path integration to remembering the location of the stool (i.e. by disorientating participants between learning/placement stages, using different starting locations for learning/placement stages and using desktop-VR with keyboard/mouse controlled movement), we included these walking-based models given the availability of body-based self-motion cues to participants during the HMD-VR and Physical environments here.

**Figure 1.**
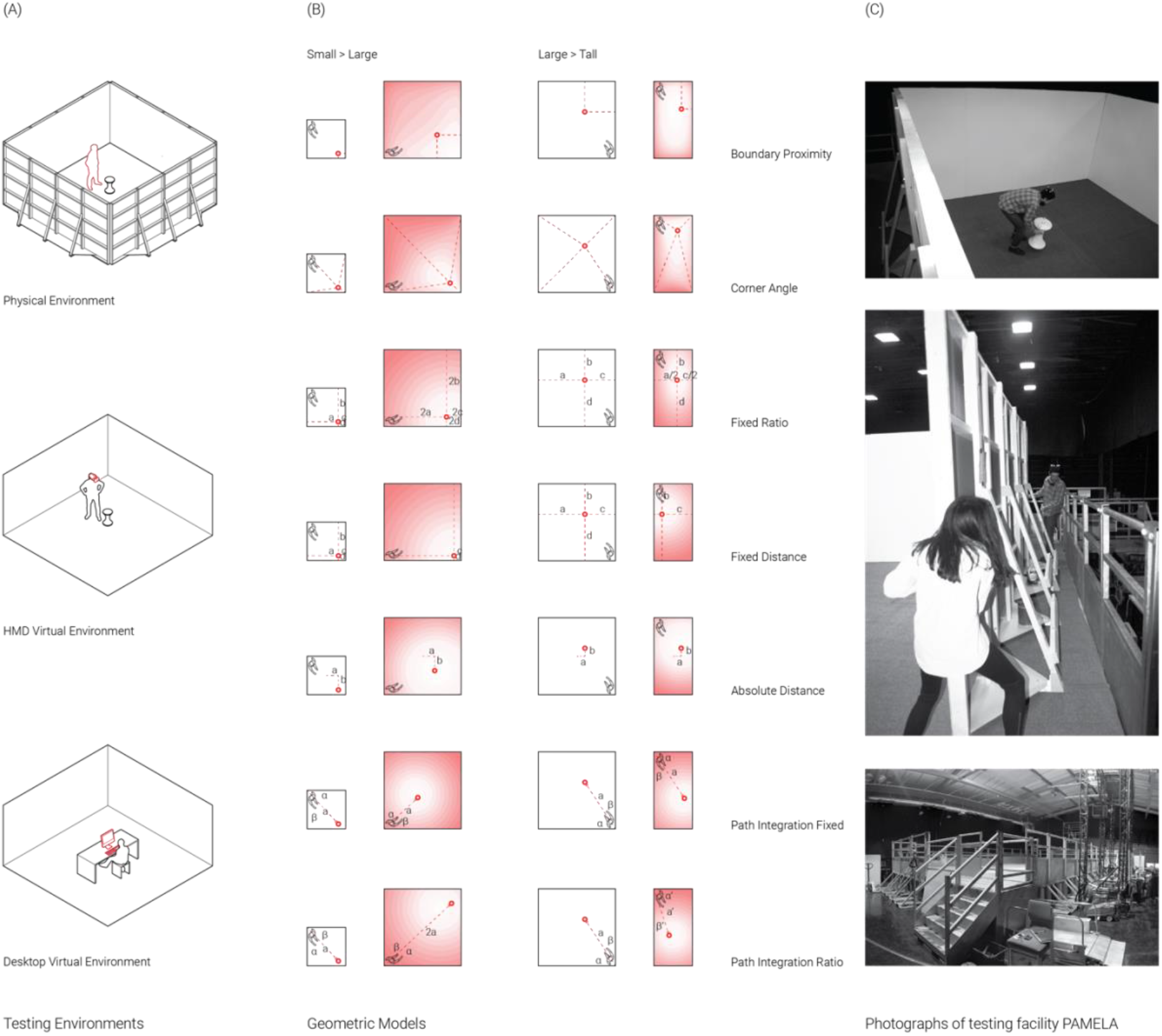
Apparatus, Procedure and Model Predictions. (A) Testing environments, top to bottom: Physical Environment; Immersive Virtual Environment; Non-Immersive Desktop Virtual Environment. (B) Experimental Trials (Small to Large, Large to Tall) and predicitve Geometric/Path Integration Models (definitions in Table 2). Each red circle corresponds to the global maximum of a model’s distribution. White shades correspond to the most likely locations, red shades to the least likely locations. Human figures indicate the corner participants entered the room and the direction they faced. (C) Photographs in the physical testing environment: Participant retrieving the object in the learning stage; Experimenters moving walls between trials; Testing facilities PAMELA platform.

**Table 2.** Definitions of predictive spatial models Note: Models BoProx, CorAng, FixRat, FixDist, AbsDist as in Hartley et al. (2004). Models PathIntFix, PathIntRat defined by authors in current study.

| Model | Definition |
| --- | --- |
| Boundary Proximity (BoProx) | Object location is represented as the 'proximity' to each of the four walls calculated by $1/(d+c)$, whereby $d$ stands for the distance to a wall, $c$ is a constant. The boundary proximity model predicts responses to tend to maintain fixed distances to closer walls (shorter distances are weighted higher in the representation) and maintain a fixed ratio for a reference in the centre of the room (opposing walls are equally weighted) |
| Corner Angle (CorAng) | Object location is represented by directions from the object to the corners of the room. The corner angle model predicts responses to be close to a location where the angles to the room's corners are as similar as possible to the angles between object and corners at initial presentation |
| Fixed Ratio (FixRat) | Object location is represented as a fixed ratio of perpendicular distances between opposite walls |
| Fixed Distance (FixDist) | Object location is represented as fixed perpendicular distance the two nearest walls |
| Absolute Distance (AbsDist) | Object location is represented as the absolute distances to all four walls. Any spatial transformation results in no one location where all four distances remain preserved |
| Path Integration Fixed (PathIntFix) | Object location is represented as the cumulative vector from trajectory from a starting location to the object. Upon spatial change, the original trajectory and vector are recreated, while maintaining facing direction |
| Path Integration Ratio (PathIntRat) | Object location is represented as the cumulative vector from trajectory from a starting location to the object. Upon spatial change, the original trajectory and vector are recreated, while maintaining facing direction and bidirectionally scaling the vector in relation to the deformed ratio of the environmental boundaries |

In Trial 1, uniform changes are made both horizontally and vertically, resulting in a displacement of the Gaussian distribution. In Trial 2 non-uniform changes are only made horizontally, resulting in a displacement and a compression of the Gaussian distribution along its horizontal axis.

### (2.5) Statistical Analyses

To assess which model best fit participants’ replacement distributions in a similar manner to Hartley et al., we first converted each model distribution into a probability density function (pdf) by normalising it so the sum of its pixels equals one. We defined the Fit Value (FV) as the value of the pdf at the position of each placement. The FV of placement (x,y) for model M is FV = M_pdf_(x,y). We then computed an average FV for each model and each environment (see Figure 2a and 2b, column D). The higher the FV, the better fit the model.

We ran two sets of analyses; in the first, we used raw participant replacement distributions. However, upon visual inspection of the raw data, we noted a high degree of disorientation in which some participants appeared to replace the object relative to the wrong corner, i.e. applying geometric principles such as fixed distances or corner angles to opposite or adjacent walls of the room to where the object was originally found (see Section 3.2). These raw placements were scattered across the four quadrants of the Trial 1 test room and across the two halves of the Trial 2 test room, resulting in FVs with large variances that rendered statistical analyses of the raw placement data less appropriate. Due to the symmetry of the environments and ability to reflect any coordinate about the centre of the room, we decided to fold replacement distributions into the ‘correctly’ orientated stool quadrant or half to simulate orientated responses (see Figure 2a). Hartley et al. 2004 performed a similar standardisation procedure to account for different cue locations across participants. However, because walking paths to learn the location crossed quadrants, path integration models (where placement was based on recalling the path walked in learning) were not valid for the folded data and so were only considered for the raw data. Given much better agreement of the folded replacement distributions with geometric model predictions, we ran a second set of analyses on these folded data for the main statistical results in Sections 3.1. For Section 3.2 and 3.3 we present analysis of the unfolded data using correlations between participant distributions and between participant distributions and predictive geometric/path integration model distributions. To perform correlations, we smoothed the data using Gaussian smoothing kernel with 20 cycles/image to reduce the binary nature of the pixels.

**Figure 2a.**
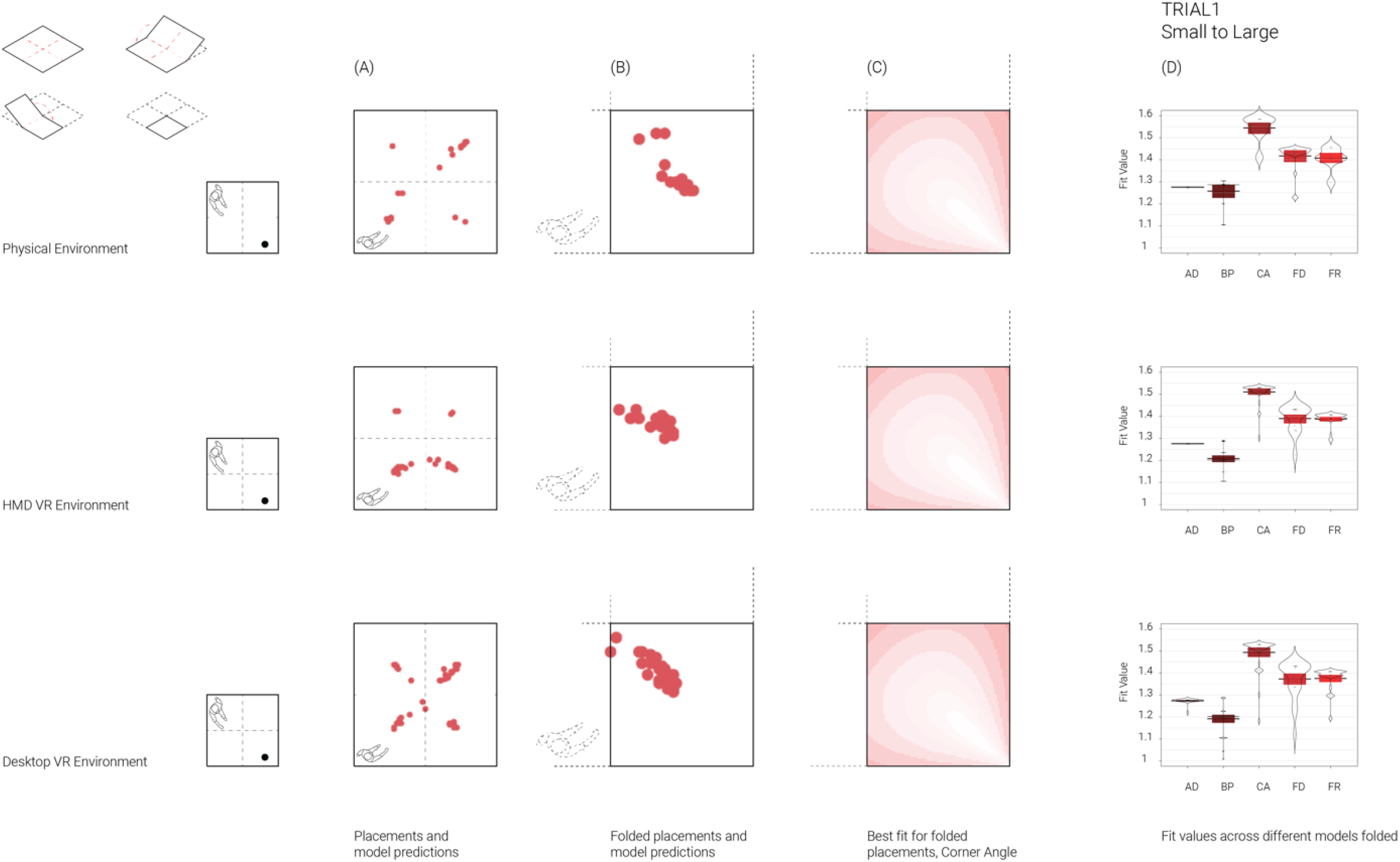
Results Trial 1, Small to Large. Top left row shows data folding operation.(A) Raw unfolded participant placements in large testing environment. (B) Placements folded into ‘correct’ south-east quadrant. (C) Best fitting Corner Angle model distribution. (D) Violin plots showing fit values for predictive models. Plots show participant FVs, the average and 95% Bayesian Highest Density Intervals. Note: because walking paths to learn the location crossed quadrants this meant that for folded data path integration models were not used for analysis of this data. AD = Absolute Distance (AbsDist,); CA = Corner Angle (CorAng); FR = Fixed Ratio (FixRat); FD = Fixed Distance (FixDist); BP = Boundary Proximity (BoProx).

In Section 3.1, for each trial using the folded data, we performed a 2-way ANOVA on FV of the placement distribution using environment (Physical, Desktop-VR and HMD-VR) as between subject factors and model as within subject factors. Bonferroni-corrected post-hoc pairwise comparisons using t-tests with pooled standard deviations were used to compare differences of between and within subject factors. Trials were also explored together using standardised FVs to account for different FV ranges (M=0, SD=1). We checked which model best accounted for participant distributions using a least square regression of the models’ pdf on the placements’ pdf, in each trial and environment, and included a weighted linear combination of all models.

## (3) Results

### (3.1) No one model explains human spatial representation

Participants were asked to memorise the location of a stool in a room and replace it after modification to the room geometry (Trial 1: small square to large square, Trial 2: large square to tall rectangle). Based on previous findings, we expected models based on fixed distances and fixed ratios of distances between the stool and the room walls best explain the replacement distributions.

For Trial 1, 86.3% of the participants’ folded placements was best accounted for by the CorAng model. We performed a mixed 2-way ANOVA on the average fit value (FV) of the 1st trial with environment as a between-subject factor and model as a within-subject factor. Both model (F(5,622) = 121.21, p<0.001) and environment (F(2,622) = 17.34, p<0.001) yielded a significant effect. Bonferroni-corrected pairwise comparisons showed that all models were significantly different from each other (all p<0.001), except FixRat and FixDist (p = 1). As shown in Figure 2a, column D, on average CorAng was the best model and BoProx the worst. Model FVs for Physical environment significantly differed from Desktop-VR (p = 0.01) but not from HMD-VR (p = 0.24), though the overall pattern of model fits was identical across all environments (Figure 2a). HMD-VR and Desktop-VR model fits did not differ from each other (p = 0.66).

In Trial 2, 62.7% of the participants’ folded placements was best accounted for by the FixRatio model. Model yielded a significant effect (F(5,634) = 46.46, p<0.001) but environment only a trend effect (F(2,634) = 2.84, p = 0.06). Pairwise comparisons showed all models were significantly different from each other (all p<0.001), except CorAng and AbsDist (p = 1). As shown in Figure 2b, column D, on average FixRat was the best model and BoProx the worst. The different environments did not significantly differ from each other in FVs (HMD-VR vs Desktop-VR: p = 1, HMD-VR vs Physical: p = 0.94, Physical vs Desktop-VR: p = 0.37).

**Figure 2b.**
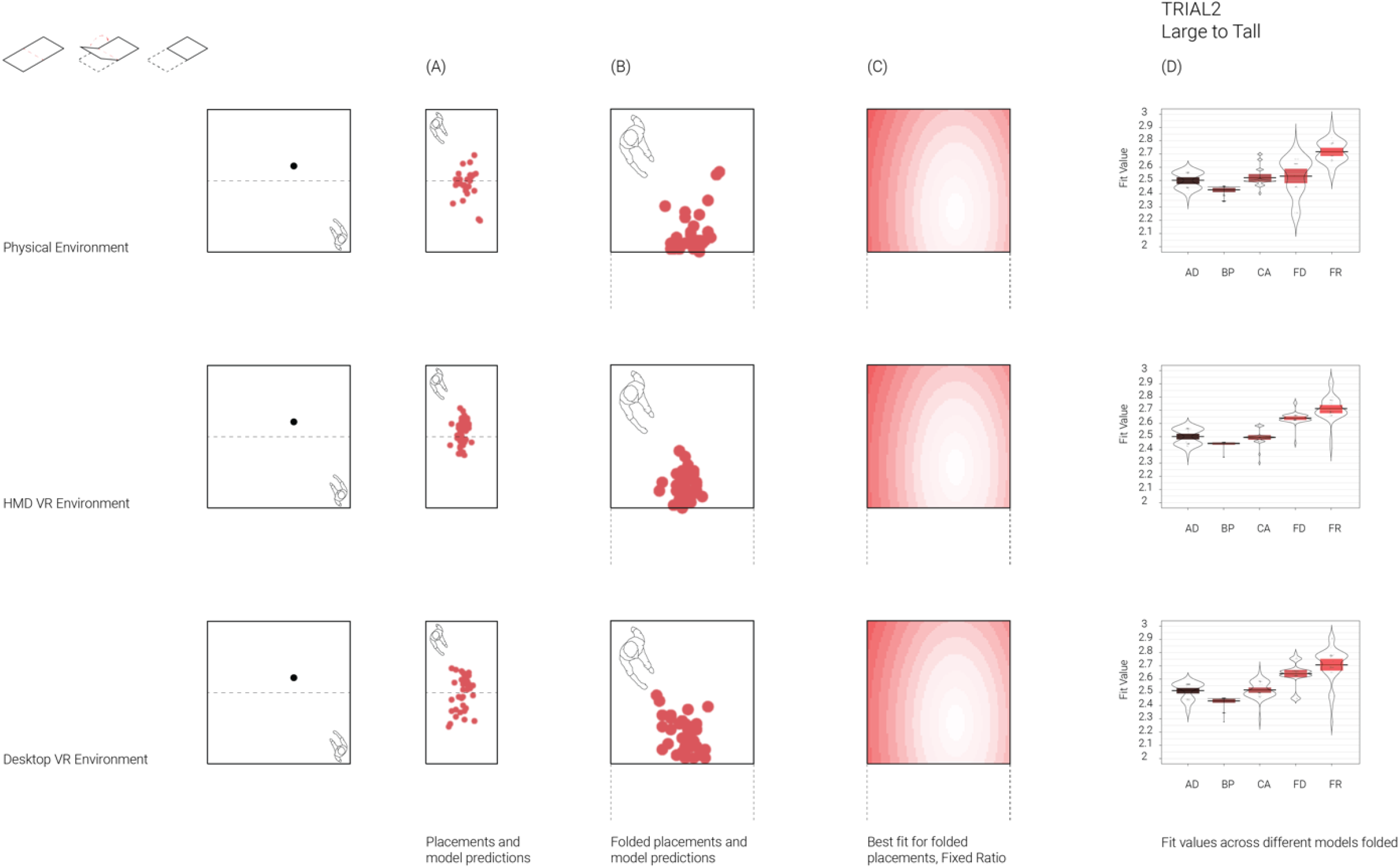
Results Trial 2, Large to Tall. Top row left shows data folding operation. (A) Raw unfolded participant placements in tall testing environment. (B) Placements folded into ‘correct’ north half. (C) Best fitting Corner Angle model distribution. (D) Violin plots showing fit values for predictive models with the average and 95% Bayesian Highest Density Intervals. Note: because walking paths to learn the location crossed quadrants this meant that for folded data path integration models were not used for analysis of this data. AD = Absolute Distance (AbsDist,); CA = Corner Angle (CorAng); FR = Fixed Ratio (FixRat); FD = Fixed Distance (FixDist); BP = Boundary Proximity (BoProx).

To examine which model best accounts for participants’ folded placements across both trial types together, we collapsed Trial 1 and Trial 2 together with z-scored FVs. This revealed FixRat best fit both trials across all environments, with other models performing similarly except for CorAng which fit better in the Physical environment (Figure 3).

**Figure 3.**
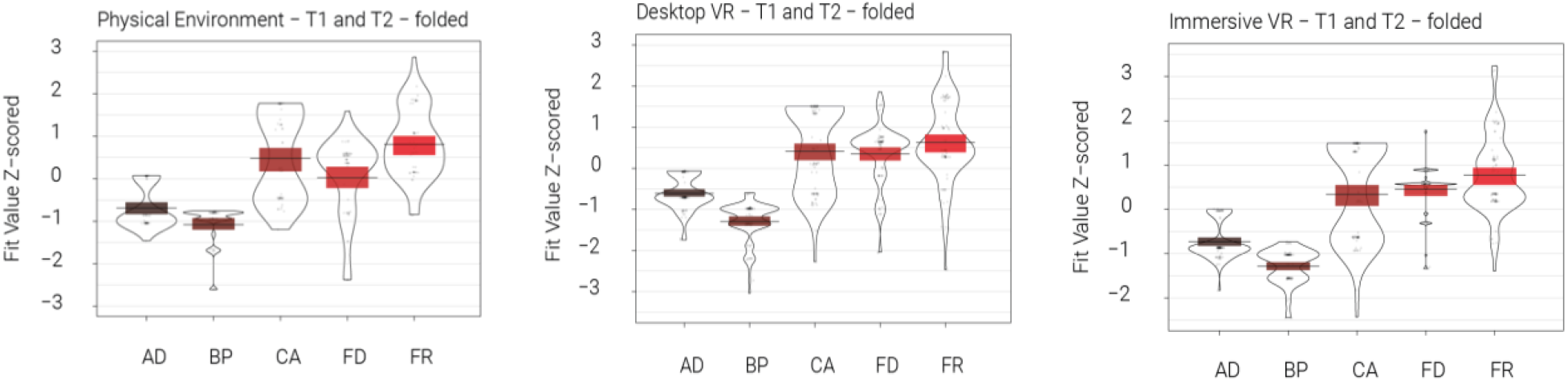
Violin plots of Z-scored fit values for predictive models on Trial 1 and Trial 2 pdf collapsed together, shown for each environment separately. Higher values indicate better model fit. AD = Absolute Distance (AbsDist,); CA = Corner Angle (CorAng); FR = Fixed Ratio (FixRat); FD = Fixed Distance (FixDist); BP = Boundary Proximity (BoProx).

Lastly, we examined the explanatory value of combining models using linear regressions. The outcome variable is the smoothed folded placements’ pdf, while the predictors are the models’ distributions. Overall, for both Trial 1 and 2 across all environments, pooling all the models’ predictions together (lin_all_models) explained more of the variance in the placements’ pdf than the best performing model’s pdf alone (CorAng in Trial 1 and FixRat in Trial 2, see Figure 4). For Trial 1 in the Physical environment, lin_all_models accounted for 2.8% of the variance in participants’ placements while CorAng accounted for 1.8%; in Desktop-VR lin_all_models accounted for 4.9% of the variance while CorAng accounted for 2.3%; in HMD-VR lin_all_models accounted for 3.2% of the variance while CorAng accounted for 1.5%. For Trial 2 in the Physical environment, lin_all_models accounted for 12.7% of the variance in the placements’ pdf while FixRat accounted for 7.9%; in Desktop-VR lin_all_models accounted for 14.4% while FixRat accounted for 7.8%; in HMD-VR lin_all_models accounted for 13.3% while FixRat accounted for 6.9%.

**Figure 4.**
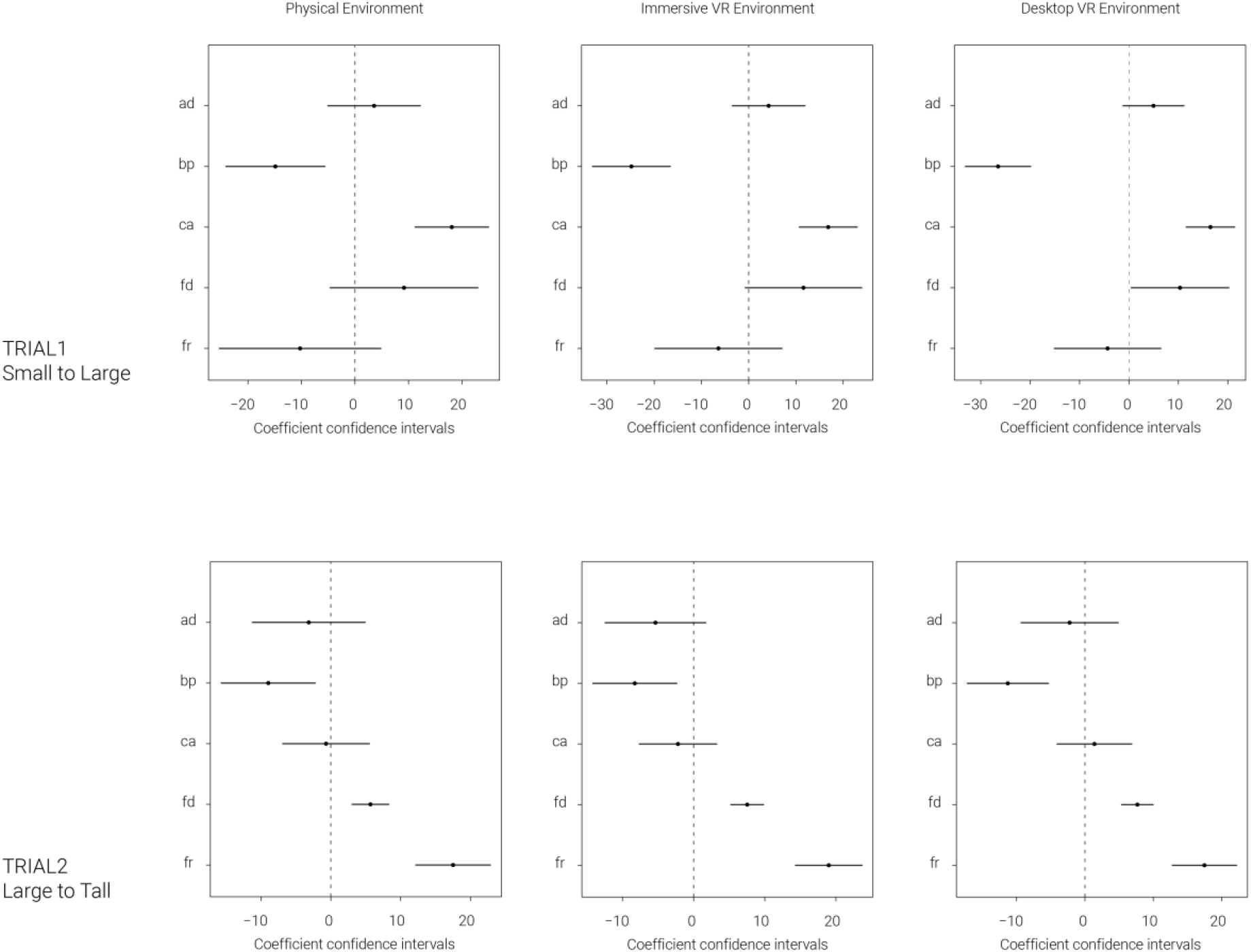
Coefficients from Least Square Regression of models’ pdf on smoothed and folded participant placements’ pdf for each trial and environment. The higher the value, the better the model explains participant placements. Negative values mean that participants tend not to place the object on the locations predicted by the model. Error bars represent 95% confidence intervals. AD = Absolute Distance (AbsDist,); CA = Corner Angle (CorAng); FR = Fixed Ratio (FixRat); FD = Fixed Distance (FixDist); BP = Boundary Proximity (BoProx).

### (3.2) HMD-VR and Desktop-VR spatial distributions of responses are more similar to each other than to Physical

Given Physical and HMD-VR environments have shared availability of walking, but HMD-VR and Desktop-VR have a shared virtual experience, we asked how their model fits and placement distributions would compare. Results from the folded data above showed identical model fit regression coefficient confidence intervals across all environments (Figure 4). Model fits significantly differed only between Physical and Desktop-VR in Trial 1 (Section 3.1), suggesting largely similar strategies across environments.

To further explore consistency between environments beyond model fits, we correlated the raw, unfolded spatial distributions of participant responses (pdf values per pixel) between environments. All environment distributions were significantly correlated with each in both Trial One and Two (Table 3, p<0.001). While correlation coefficients were generally higher for Trial 2 (which also showed better R^2^ model fits than Trial 1 above), the two VR environments showed higher coefficients across both trials (r = 0.90 and r=0.79, respectively).

**Table 3.** Correlation coefficients between raw, unfolded participants’ placement distributions in the three types of setup in (a)Trial 1 and (b)Trial 2.

| **Trial 1** | Physical | Desktop VR | HMD VR |
| --- | --- | --- | --- |
| Physical | 1 | 0.20 | 0.18 |
| Desktop VR |  | 1 | 0.90 |
| HMD VR |  |  | 1 |

| **Trial 2** | Physical | Desktop VR | HMD VR |
| --- | --- | --- | --- |
| Physical | 1 | 0.57 | 0.65 |
| Desktop VR |  | 1 | 0.79 |
| HMD VR |  |  | 1 |

To better understand the smaller correlation coefficients between Physical and both VR environments in Trial One, we examined if the precision of distance estimation contributed to differences between them, noting on visual inspection of both the raw and folded data that the Physical distributions appeared more proximal to the walls than in the VR. By measuring the distance between the pixel with the highest density of placements and the nearest two walls per environment in the folded data of both trials, we found indeed that Physical placements were on average closer to the walls than other environments in Trial 1 but not Trial 2 (Table 4).

**Table 4.**
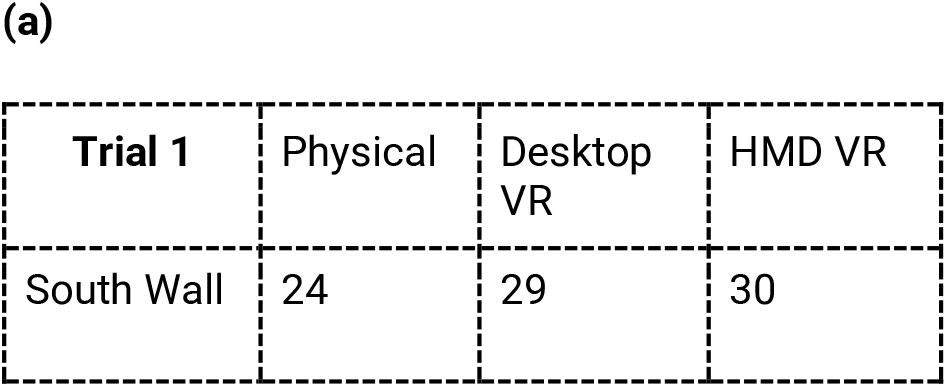
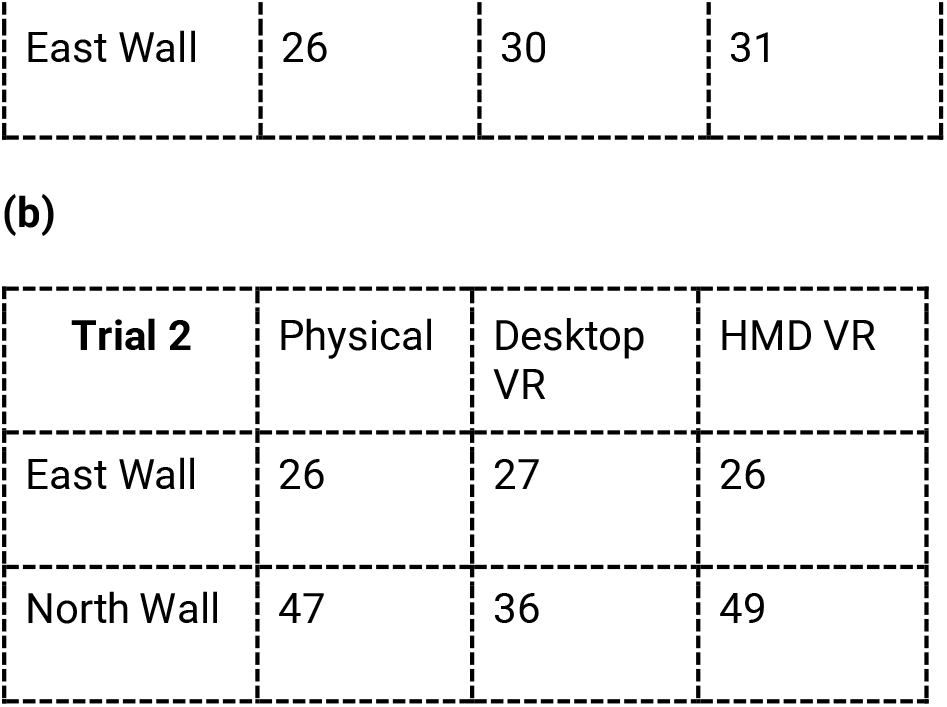
Distances (number of pixels) between the pixel with the highest density of replacements and the two nearest walls in (a) trial 1 and (b) trial 2 using folded data. Walls labelled by cardinal directions with north taken as the top wall in Figures 2a and 2b.

### (3.3) Path integration contributions in the unfolded data

Finally, we returned to the raw, unfolded data to understand the impact of path integration across environments. Despite very poor fit to the models in the raw data we aimed to assess the influence of walking in HMD-VR and Physical for completeness of comparing environments. Using correlations, we explored how path integration (PI) and geometric based model distributions compared to the raw unfolded data. Across all environments, PathIntFix best correlated with the raw placement distributions for Trial 1. For Trial 2, both PI models correlated best for Physical and second best for the VR environments behind the BoProx model (Figure 5).

**Figure 5.**
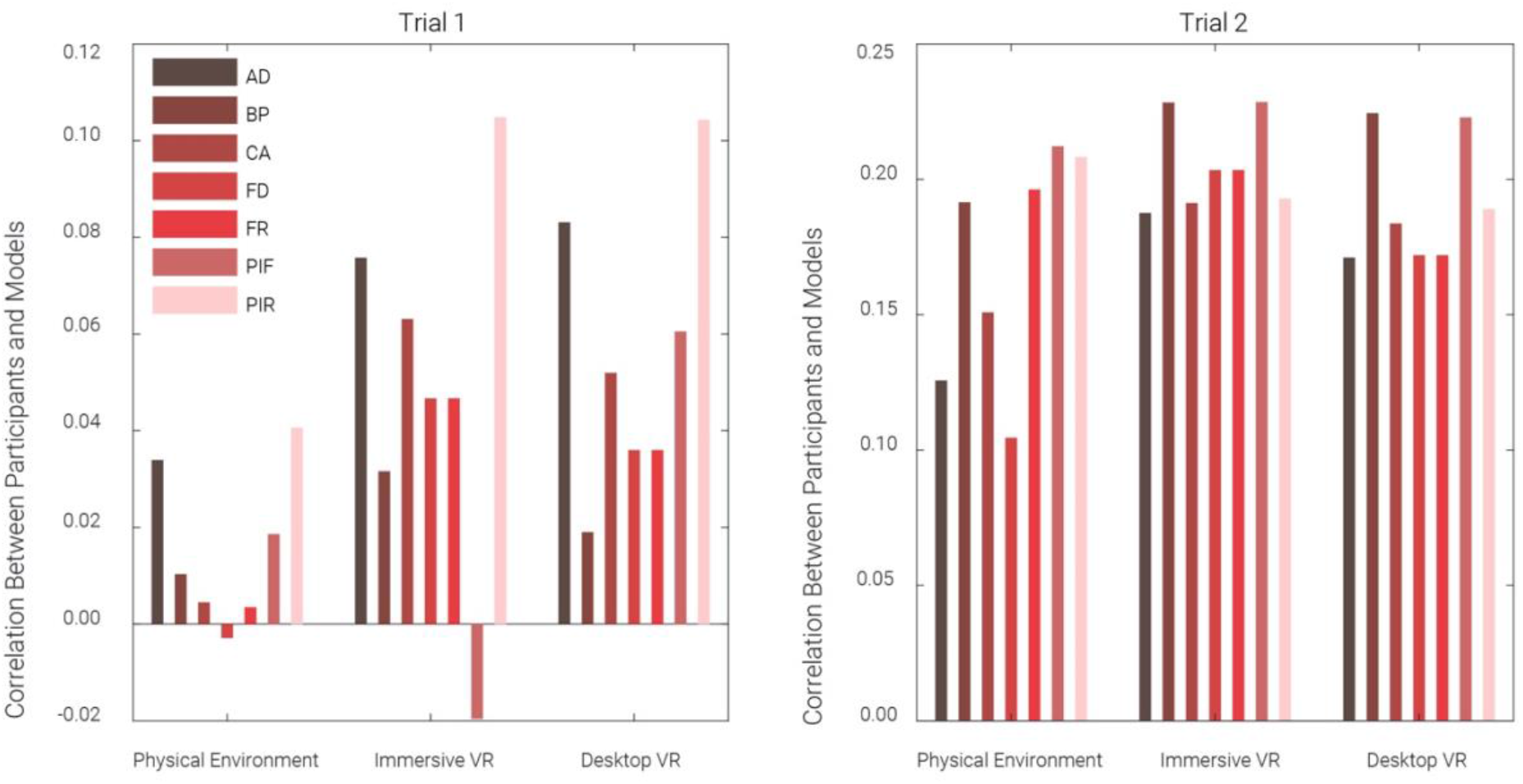
Correlations between distributions of participant group responses and geometric/path integration models for raw unfolded data across environments. PIF = Path Integration Fixed (PathIntFix); PIR = Path Integration Ratio (PathIntRat); AD = Absolute Distance (AbsDist,); CA = Corner Angle (CorAng); FR = Fixed Ratio (FixRat); FD = Fixed Distance (FixDist); BP = Boundary Proximity (BoProx).

The comparable correlation coefficients of PI based models in the Desktop-VR and Physical environments suggests no strong influence of body-based self-motion cues to performance. However, Physical environment coefficients for all models were generally poorer than for both VR environments, especially for Trial 1. To examine this in greater detail, we assessed levels of disorientation and replacement variance across environments and trials. Visual inspection of the raw data in Trial 1 shows four clusters of responses in each room quadrant, suggesting application of geometric principles to the wrong axes of the room and thus disorientation of some participants relative to the amber lighting cue. We quantified the proportion of responses in the ‘correct’ quadrant per environment - the quadrant of the room in which the stool was originally found (Figures 2A and 2B) - which revealed greater disorientation for the Physical environment, where only 8.7% of responses were seemingly orientated (Table 5A). Furthermore, we found that the Physical Trial 1 has the greatest variance of all environments and trials, with the lowest consistency between individuals in estimating the location of the stool (Table 5B).

**Table 5.**
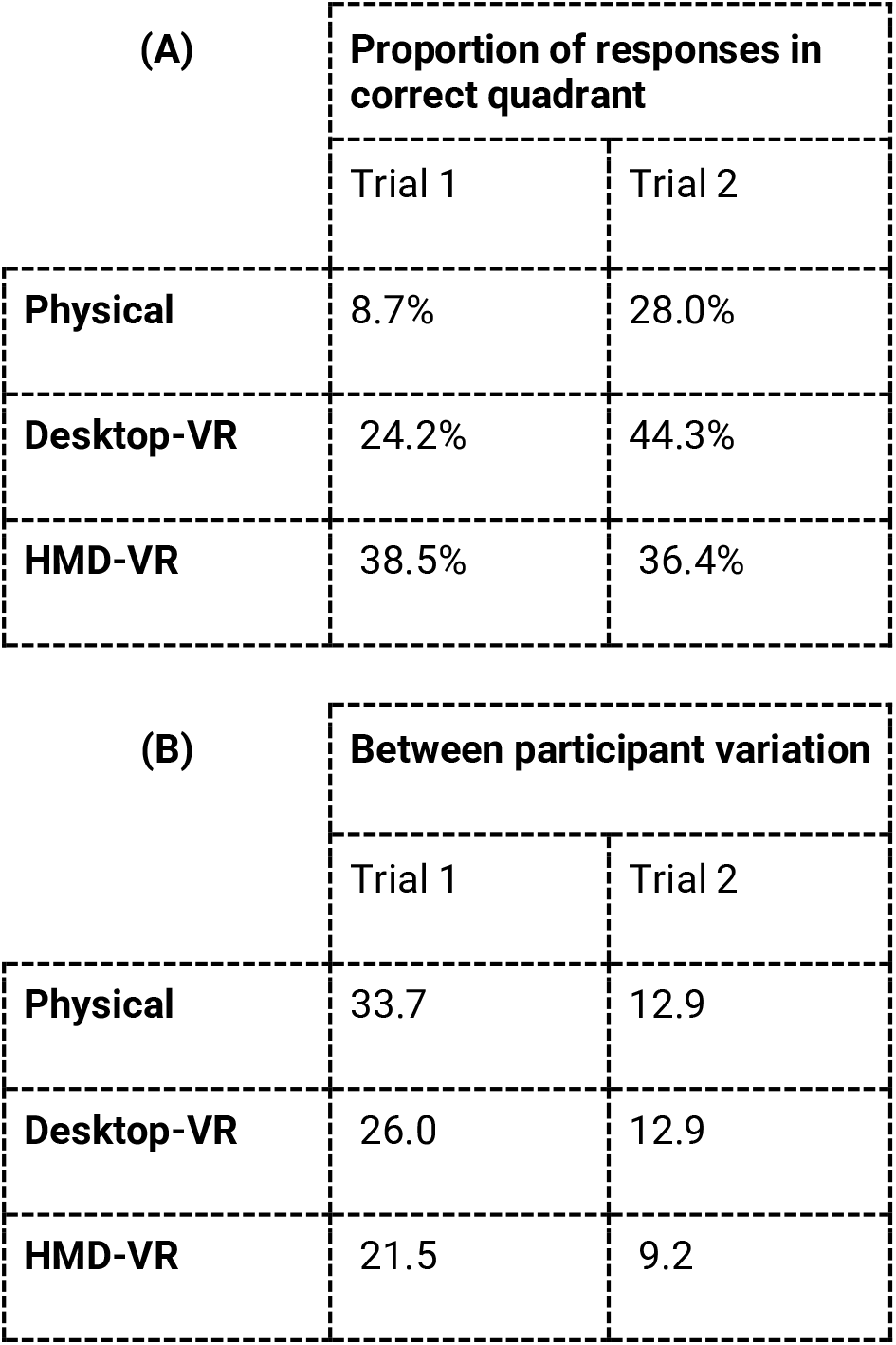
Disorientation and consistency between results per trial across environments for raw data. A. Percentage proportion of responses found in correct environment quadrant (Trial 1) or half (Trial 2) i.e. where stool was found in learning stage. B. Variation in responses between participants quantified using the average distance in pixels between randomly paired response locations of each participant separately in each environment and trial type. Higher values indicate increased distance between participant locations and therefore higher variation.

| (A) | Proportion of responses in correct quadrant |  |
| --- | --- | --- |
|  | Trial 1 | Trial 2 |
| Physical | 8.7% | 28.0% |
| Desktop-VR | 24.2% | 44.3% |
| HMD-VR | 38.5% | 36.4% |

**Table 5.** Disorientation and consistency between results per trial across environments for raw data.
| (B) | Between participant variation |  |
| --- | --- | --- |
|  | Trial 1 | Trial 2 |
| Physical | 33.7 | 12.9 |
| Desktop-VR | 26.0 | 12.9 |
| HMD-VR | 21.5 | 9.2 |

## (4) Discussion

We had participants learn the location of a stool in a room and subsequently replace it after we manipulated the boundary size or geometry. We repeated this in a Physical, Desktop-VR and head mounted display (HMD) immersive-VR room and compared participant response distributions to seven different models of spatial representation. Based on previous findings relating human spatial memory to a model of place cell firing (Hartley et al., 2004), we predicted participants would use a model of fixed distances between the stool and the two nearest walls to complete Trial 1, and a model of fixed ratios of distances between the object and all four walls for Trial 2.

Firstly, results showed that indeed a Fixed Ratio model best fit response distributions for Trial 2, but response distributions for Trial 1 were better accounted for by a Corner Angle model using the angles subtended between the stool and the four room corners. Secondly, this pattern of model fits was identical across Physical, Desktop-VR and HMD-VR environments. Finally, closer examination of the response distributions themselves showed varying rates of disorientation and between subject consistency across the two transformation types. Thus, while different small-scale boundary manipulations differentially influence human spatial memory, this influence is similar across virtual and physical environments.

### (4.1) Replications of Hartley et al (2004)

Our findings partly replicate Hartley et al. (2004) on several levels. Fixed Ratio and Corner Angle models performed second and third best respectively in their Desktop-VR only study. Their similarly good fit here across Desktop-VR, HMD-VR and Physical environments underscores the dependence of human spatial representation on distance and proportion information. These specific models better capture the shape of responses distributions by allowing for stretching or skewing of responses, unlike Absolute Distance or Fixed Distance which are symmetrical by nature (Hartley et al., 2004).

Fixed Ratio is a model inspired by the influence of boundaries on place cell firing. By replicating the Fixed Ratio model’s predictive power not just for Trial 2 but also for both Trials collapsed together across the three environments corroborates the idea that aspects of spatial behaviour may relate to the neural dynamics in the hippocampal region. Indeed recently, a HMD-VR study also similar to Hartley et al. showed distortions in participant object replacements following boundary manipulations were dependent on which boundary was most recently visited, and this was identical to findings of grid cell distortions in parallel rodent studies (Keinath, Epstein, & Balasubramanian, 2018; Keinath et al., 2021). Additionally, non-orthogonal boundary distortions resulted in corresponding non-orthogonal distortions to both rodent grid cell firing and human spatial memory representations (Krupic et al., 2015)(Bellmund et al., 2020).

Another point of replication is the high level of participant disorientation seen here and in the second experiment of Hartley et al (2004). Their second experiment more closely resembled our protocol in which they also used the same learning stage stool position across participants. Comparing experiment two trials with similar transformations and stool positions to Trial 1 and 2 here reveals remarkably similar response distributions to our unfolded data; four clusters of disorientated replacements for Trial 1 and an elongation of replacements for Trial 2 (Figure 6). This is notable considering their more salient mountain orientation cue. Boundaries and geometry may therefore be as critical to orientation as specific directional cues (Cheng & Newcombe, 2005), which originally was shown in children (Hermer & Spelke, 1994) and more recently explored in adults (Kelly et al., 2009). This may explain the variation in participant consistency and disorientation between Trial 1 and Trial 2, in which Trial 1 involves a symmetrical square test room but Trial 2 a more geometrically informative polarised rectangular environment.

**Figure 6.**
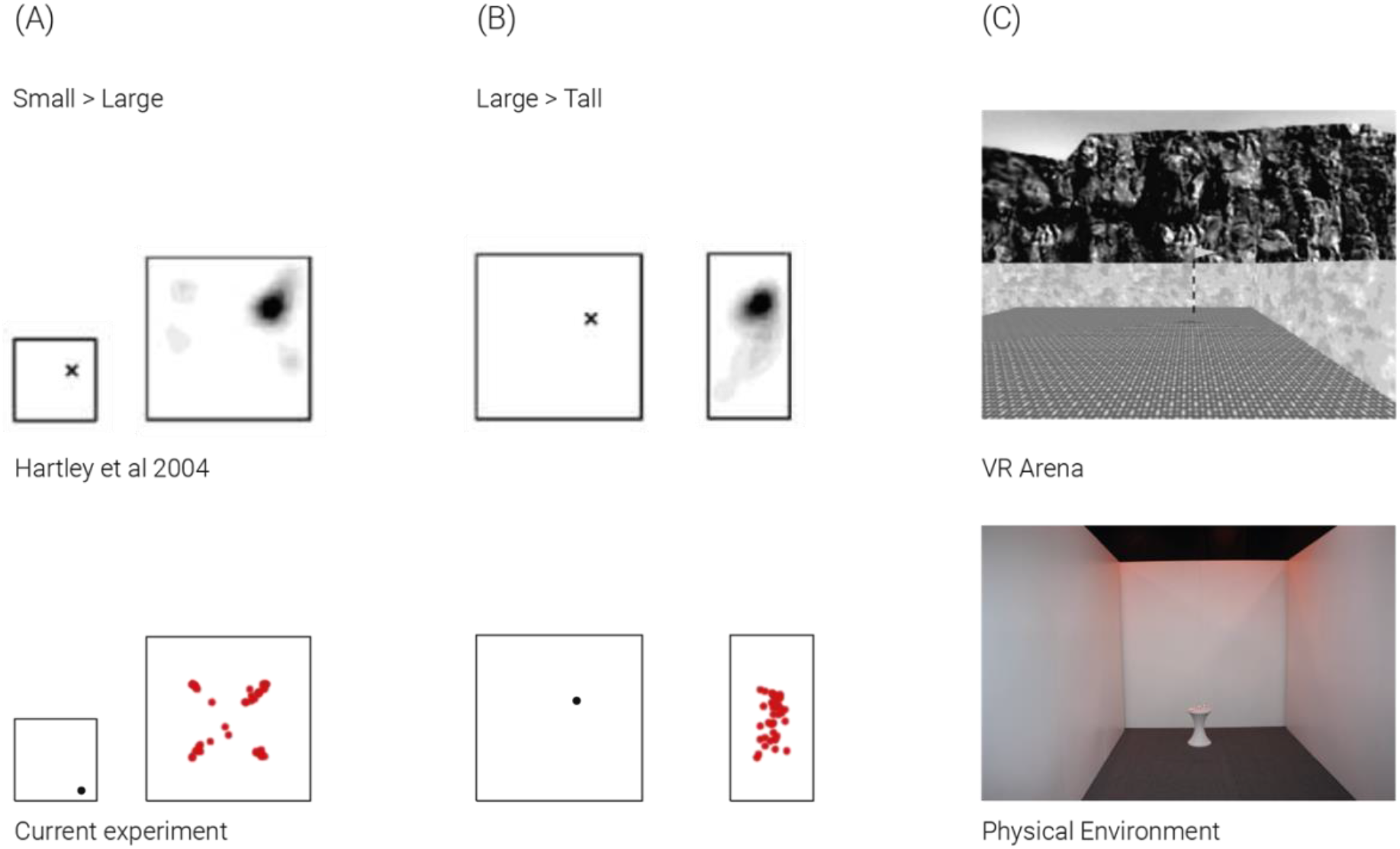
Comparison of results between Hartley et al. 2004 experiment two in desktop VR (top panel) and results here in desktop VR environment (bottom panel) for Trial 1 and Trial 2 with similar transformations and stool positions. Photos on far right show differences in orientation cue saliency: the virtual environment of Hartley et al. 2004 (top) with mountains projected at infinity along one wall, and physical environment here (bottom) that was laser scanned into virtual reality with orange lighting cue along one wall.

Some differences to Hartley et al (2004) were found here. Principally, we found a poor fit of their overall Boundary Proximity model (of which Fixed Ratio is a component) and best overall fit of weighted linear combination of all models which was not examined in their study. One possibility for the poor fit of the Boundary Proximity model here could be limitations in our protocol. Here, we (A) used the same learning stage stool location across participants, (B) we were limited to two transformation types and (C) had high levels of participant disorientation necessitating that we fold the response distributions for analysis. These factors may disrupt the effectiveness of the Boundary Proximity model, which was able to predict behavioural responses based on the interaction of multiple stool placements and transformation types. We were unable to use more transformation types or stool placements owing to the study design with the Physical environment; the first iteration in the Physical environment in PAMELA involved manual moving of walls which was time and energy intensive, making it difficult to repeat multiple trial types per participant. Unfortunately, Hartley et al. (2004) did not fit the Boundary Proximity model to their experiment two data, which also has the same stool placement across participants.

The best fit of the weighted linear combination of all spatial models potentially reflects the larger inter-individual variation in determinants and strategies in human spatial representation (Hegarty & Waller, 2005); Coutrot et al., 2018; Spiers, Coutrot, & Hornberger, 2021). Indeed, one advantage of our study over Hartley et al (2004) is the much larger and more heterogeneous study population in terms of age and gender. Unfortunately, the sample sizes did not permit us to explore differences in model fits across these sub-populations.

### (4.2) Contribution of path integration

We aimed to expand on the Desktop-VR based findings of Hartley et al (2004) by exploring the impact of path integration to remembering the location of the stool. We included both a Physical and HMD-VR replication of the Desktop-VR task where participants had full access to natural walking rather than mouse and keyboard-controlled motion. Previous findings show path integration based distance estimates are influenced by boundary expansion and contraction (Chen, He, Kelly, Fiete, & McNamara, 2015). Our path integration models made predictions that the stool should be replaced across quadrants from the starting location, which are only interpretable in the unfolded raw data. Because the raw data patterns implied participants were disoriented with regard to direction, we largely examined the folded data which precluded the path integration models. While the path integration models showed a higher correlation than other models with the stool placement patterns in the raw data, these correlations were relatively low. It is possible that with some disorientation in an ambiguous environment, participants defer to memory of previous paths to remember locations - as observed in a study that unexpectedly removed environmental landmarks during a homing task (Zhao & Warren, 2015), and another study that put visual cues in conflict with path integration (Sjolund, Kelly, & Mcnamara, 2018). Future research with clear orientation cues and a larger environment to increase the discrepancy between path integration models and geometric cues would be helpful in exploring this.

### (4.3) Consistency across physical and virtual environments

We found a remarkable degree of concordance for model fits of response distributions across physical, desktop-VR and HMD-VR, suggesting similar use of distance and proportion information to perform the task despite differences in availability of body-based cue information. However, when comparing the response distributions themselves, we found subtle differences in distance estimation and disorientation. Physical responses were slightly closer to the walls than in the VR experiments, replicating known underestimation of distances due to differences in depth perception in virtual environments (Jones, Swan, Singh, Kolstad, & Ellis, 2008; Wann, Rushton, & Mon-Williams, 1995). Despite this, all environment responses conserved the relative distances between the two walls, revealing unperturbed estimates of proportion.

The greater participant variation in the Physical environment, especially with the more ambiguous Trial 1 geometry, is likely due to greater participant disorientation, as reflected in very low proportions of replacements in the correct Physical room quadrant. One explanation for lower rates of VR disorientation is the novelty of interacting with VR environments (this study was run in 2015). This may have prompted participants to look up more to fully explore their environment and therefore become more aware of the lighting cue, but we cannot confirm this without heading angle tracking. However, these findings replicate a separate study showing that, when geometric and featural cues were in conflict, participants in HMD-VR were more likely to weight the featural cue over geometry than in the physical version (Kimura et al., 2017).

In sum, findings here support the general use of VR for small-scale spatial memory tasks. This may also extend more broadly. Recent evidence indicates that similar brain activation patterns occur during spatial memory retrieval of a large environment when either immersive-VR with body based cues was used or not (Huffman & Ekstrom, 2019). Despite this convergence it is still important to consider that neuropsychological or architectural research studies may opt for HMD-VR, due to purported higher degree of environment presence due to access to physical motion with less sensory conflict (Weech, Kenny, & Barnett-Cowan, 2019). HMD-VR may be particularly useful in looking at head-scanning behavior in future studies. The VR experiments in this current study were easier and less time-consuming to run than the Physical experiment. Besides known variations in depth perception, results suggest minimal variation in strategy use between environments with different sources of spatial information. However, findings from more complex and large-scale wayfinding tasks are conflicting, and often depend on the task goal and outcome metrics (Sousa Santos et al., 2009; Srivastava, Rimzhim, Vijay, Singh, & Chandra, 2019). Recently, a novel study compared HMD and Desktop-VR wayfinding on a battery of different tasks in a multi-story building with different rooms. Findings showed Desktop-VR users looked around less, travelled faster and with shorter distances (Feng et al., 2022). However, overall route choice was similar, and more critically, user experience in perceived presence, usability, cybersickness or realism was identical between the two groups. Such large-scale studies with multiple task components will be key to mapping consistent differences between physical, desktop and immersive virtual environments, such as depth perception, travel time / distance estimates (Brunec, Javadi, Zisch, & Spiers, 2017) and map drawing (Jafarpour & Spiers, 2017) which will enable informed future choices between environments for a given task.

## (5) Concluding remarks

This study aimed to investigate how manipulation of wall boundaries impacts human spatial memory of an object’s location, and how this compares across physical and virtual environments. We found different boundary manipulations vary in how they affect human spatial memory in a small-scale room. Unlike previous findings upon which this study is based, results here showed no single predictive geometric model of spatial representation could best predict responses. However, we do replicate findings that models derived from place cell firing can partially predict these responses and expand these to show this is consistent in Desktop-VR, HMD-VR and physical environments, despite differences in body-based cue availability. Overall, we show that Desktop and HMD-VR allow a good and interchangeable approximation for examining the varying impact of boundaries on human spatial memory in small-scale physical environments.

## (6) Acknowledgments

The authors wish to thank Nikos Papadosifos, Derrick Boampong, Tatsuto Suzuki, Biao Yang, Aaron Breuer-Weil, Christopher Crispin, Simon Julier, Sebastian Friston, Panos Mavros, Dominik Zisch, Charles Middleton, Rowan Haslam, Ludovico Saint Amour Di Chanaz and Thomas Reed. This work was supported by a UCL GRAND CHALLENGES GRANT and a James S. McDonnell Foundation Scholar grant awarded to Hugo Spiers. Declarations of interest: none.

## Author contributions

**Fiona E. Zisch:** investigation, writing – original draft, project administration, visualisation. **Coco Newton:** investigation, writing – original draft/review & editing, formal analysis, project administration. **Antoine Coutrot:** formal analysis, data curation. **Maria Murcia:** investigation, software. **Anisa Motala:** investigation. **Jacob Greaves:** investigation. **William de Cothi:** investigation, software. **Anthony Steed:** resources. **Nick Tyler:** resources. **Stephen A. Gage:** resources. **Hugo J. Spiers:** conceptualisation, methodology, Writing – review & editing, funding acquisition, supervision

## Notes

### Competing Interest Statement

The authors have declared no competing interest.

